# Quantitative proteomic alterations of human iPSC-based neuronal development indicate early onset of Rett syndrome

**DOI:** 10.1101/603647

**Authors:** Suzy Varderidou-Minasian, Lisa Hinz, Dominique Hagemans, Danielle Posthuma, Maarten Altelaar, Vivi M. Heine

**Affiliations:** Biomolecular Mass Spectrometry and Proteomics, Bijvoet Center for Biomolecular Research and Utrecht Institute for Pharmaceutical Sciences, University of Utrecht, Padualaan 8, 3584 CH Utrecht, the Netherlands; Netherlands Proteomics Center, Padualaan 8, 3584 CH Utrecht, the Netherlands; Department of Complex Trait Genetics, Center for Neurogenomics and Cognitive Research, Amsterdam Neuroscience, Vrije Universiteit Amsterdam, the Netherlands; Clinical Genetics, Amsterdam UMC, Amsterdam Neuroscience, Vrije Universiteit Amsterdam, Amsterdam, the Netherlands; Pediatric Neurology, Emma Children’s Hospital, Amsterdam UMC, Amsterdam Neuroscience, Vrije Universiteit Amsterdam, Amsterdam, the Netherlands..

**Author notes:** These authors contributed equally to this work (Suzy Varderidou-Minasian, Lisa Hinz, Maarten Altelaar, Vivi M. Heine).

**Keywords:** Rett syndrome, iPSC, neuron differentiation, quantitative mass spectrometry, TMT-10plex

## Abstract

Rett syndrome (RTT) is a progressive neurodevelopmental disease often caused by mutations in the X-linked gene encoding methyl-CpG binding protein 2 (MeCP2). The mechanisms by which impaired MeCP2 induces the pathological abnormalities in the brain are not understood. To understand the molecular mechanisms involved in disease, we used an RTT patient induced pluripotent stem cell (iPSC)-based model and applied an in-depth high-resolution quantitative mass spectrometry-based analysis during early stages of neuronal development. Our data provide evidence of proteomic alteration at developmental stages long before the phase that symptoms of RTT syndrome become apparent. Differences in expression profiles became more pronounced from early to late neural stem cell phases, although proteins involved in immunity, metabolic processes and calcium signaling were already affected at initial stages. These results can help development of new biomarkers and therapeutic approaches by selectively target the affected proteins in RTT syndrome.

## Introduction

Rett syndrome (RTT) is a severe neurodevelopmental disorder that mainly affects females with a frequency of ~1:10,000 [1]. Clinical features of RTT start presenting themselves around 6-18 months of age, and include deceleration of head growth, abnormalities in cognitive, social and motor skill development and seizures [2, 3]. Postmortem studies showed that RTT patient brains present with an increased density of neurons in combination with reduced soma sizes, compared to healthy control brains [4, 5]. RTT neurons further show a decrease in dendritic branching, and a reduced number of dendritic spines and synapses [6, 7], typical for an immature phenotype, therefore categorizing RTT as neurodevelopmental disorder [1]. The molecular mechanisms underlying the onset of neuronal development in RTT is however unknown.

In 90-95% of RTT cases, the disease is caused by a loss-of-function mutation in the X-linked gene encoding methyl-CpG binding protein 2 (MeCP2) [8]. MeCP2 mutations predominantly occur on the X chromosome, which upon random X chromosome inactivation in females results in somatic mosaics with normal and mutant MeCP2 [9]. Males carrying a MeCP2 mutation are not viable or suffer from severe symptoms and die early in life [10]. MeCP2 was first described as a nuclear protein modulating gene expression, via binding to methylated DNA and hundreds of target genes. These modulations take place through direct repression or activation of genes, or by means of DNA modulation and secondary gene regulation. Consequently, mutations in MeCP2 lead to miss-regulation of hundreds of genes, including those influencing brain development and neuronal maturation [11-14]. So far research in RTT focused on genomic and transcriptomic studies [15-17] and less so on proteome changes [18, 19]. Recent advances in mass spectrometry-based proteomics now facilitate the study of global protein expression and quantification [20]. Considering the broad and complex regulating functions of MeCP2, modulating multiple cellular processes, we need insight into the final molecular effectors reflected by perturbation at the protein level to understand pathological states.

Here we used an iPSC-based RTT model and performed proteome analysis on iPSC-derived neuronal stem cells (NES cells) [21]. Earlier studies proved that iPSCs from RTT patients reflect disease-specific characteristics, including changes in neuronal differentiation at early stages of development [22, 23]. However, we lack knowledge on the precise molecular mechanisms underlying the progression of the disease. To study early alterations in the proteome of RTT cells compared to isogenic controls (iCTR), we performed a high-resolution mass spectrometry-based quantitative proteomics at different time points during neuronal stem cell development (Fig. 1). Interestingly, proteins differentially expressed in RTT versus iCTR are involved in cellular processes, implicated in classical features of typical RTT phenotypes, such as dendrite formation and axonal growth. We show that the difference between RTT and iCTR, in terms of the number of differentially expressed proteins, already begins at early stages and increases at later neural stem cell stages. We also show that a specific group of proteins involved in immunity and metabolic processes are differentially expressed in RTT at all time points studied. Here we provide evidence of target proteins that could be used as potential targets for early treatments to reduce the progression of RTT symptoms.

**Figure 1.**
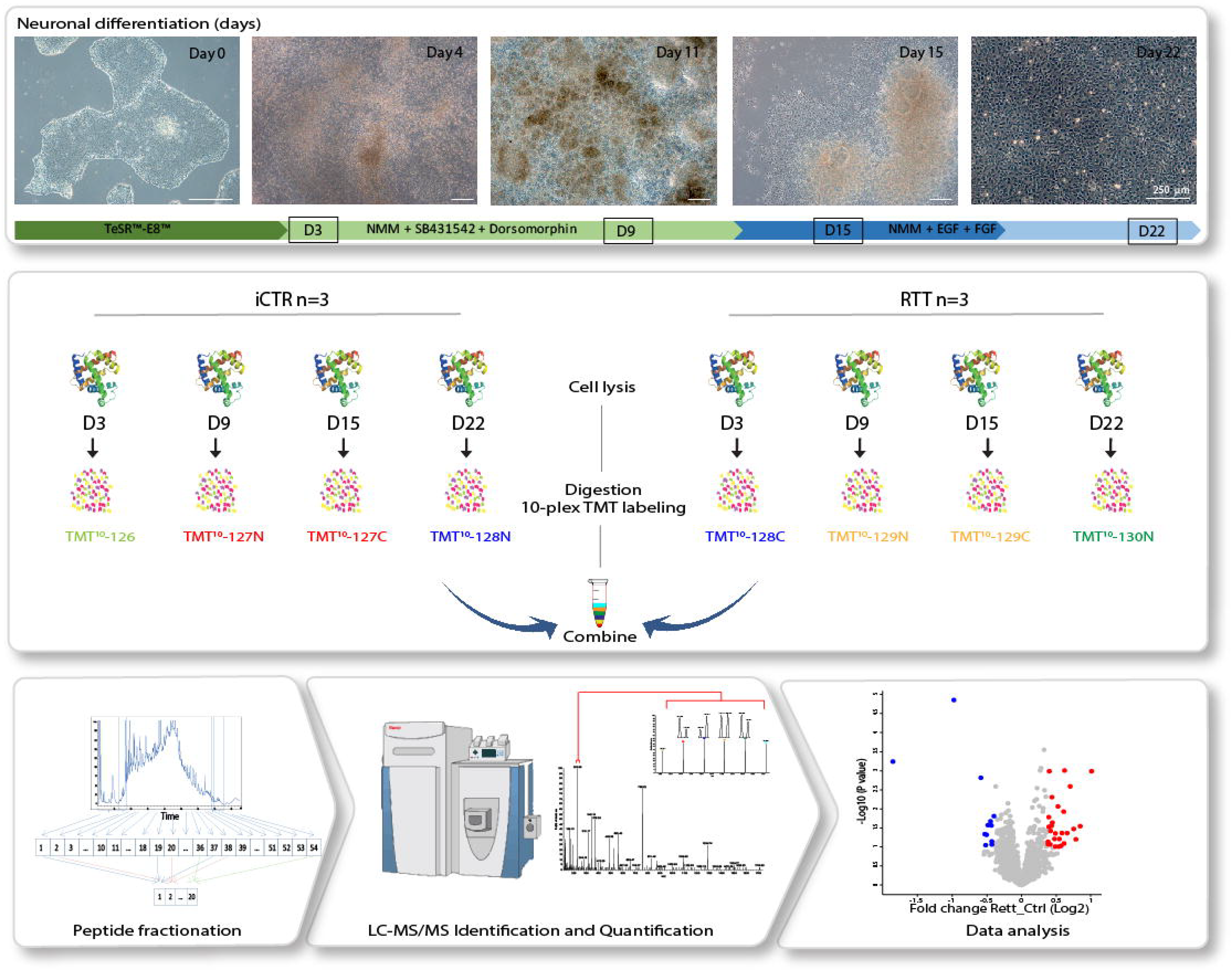
Overview of experimental workflow. iPSC differentiation towards neurons. Different colors in arrows indicate change of medium. Squares mark days of sample collection. Samples at indicated time points for iCTR and RTT were processed for proteomic analysis (n=3). In total 24 samples were subjected for tryptic digestion, TMT-based isotope labeling, high-pH fractionation and LC MS/MS analysis. Different bioinformatic approaches were then used to analyze the data.

## Results

### Generation of iPSCs from RTT and isogenic controls

RTT patient and iCTR fibroblasts were reprogrammed into iPSCs via electroporation of reprogramming plasmids as published before [24]. Pluripotency was confirmed using the PluriTest analysis (**Fig. S1C**), and if needed further validated using classic assays, i.e. immunocytochemistry (**Fig. S1A**) and RNA expression (**Fig. S1B**). Expression for normal and affected MeCP2 in the RTT and iCRT lines was confirmed by immunocytochemistry and PCR for MeCP2 (Fig. 2A, B). Additionally, the protein expression level of MeCP2 was validated by mass spectrometry over time and showed a higher expression of MeCP2 in iCTR samples relative to RTT lines (Fig. 2C). Generation of NES cells was performed as described before [21]. Neuronal induction was confirmed by morphological alterations of the cells such as neuronal rosette appearance after approximately 12 days of neuronal development initiation (Fig. 1).

**Figure 2.**
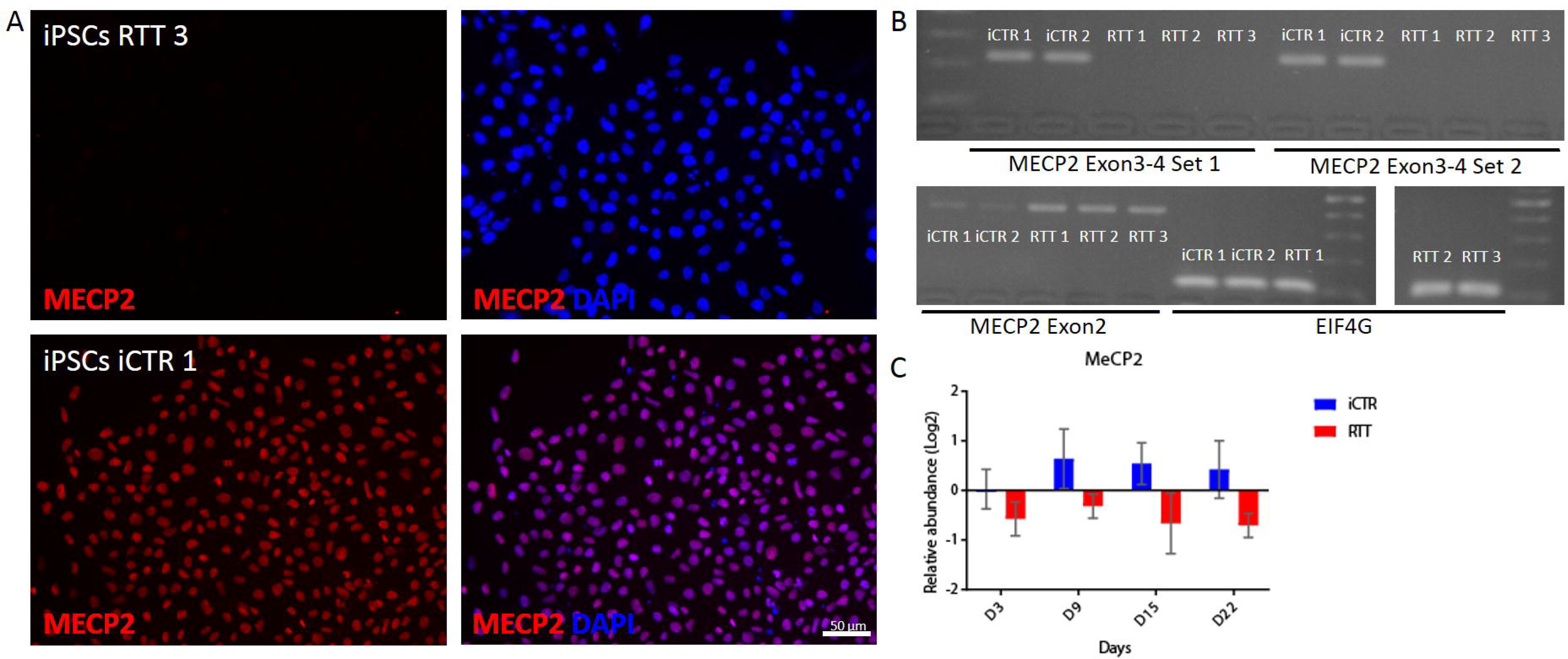
iCTR and RTT cell line validation. (A) Representative immunohistochemical staining for MeCP2 in RTT iPSC line (upper) and iCTR iPSC line (lower). (B) PCR results of iCTR and RTT iPSC lines for two different primer sets spanning deletion Del_Ex3-4 and Exon2 as positive control. (C) Relative abundance level of MECP2 in RTT NES and iCTR NES lines at different time points of sample collection.

### MS-based quantitative proteomics during neuronal development

To get an overview of the proteomic changes between RTT and iCTR during neuronal development, cell lysates at indicated time points were subjected to tryptic digestion, high-pH fractionation followed by high-resolution tandem mass spectrometry (LC-MS/MS) analysis and TMT-10plex quantification (Fig. 1). In total we identified 7720 proteins, of which 3658 proteins were quantified in all samples (**Fig. S2**). Next, to determine protein expression changes over time-points, we compared RTT versus iCTR and considered proteins with a p-value ≤ 0.1 and ≥ 1.3 fold change in 2 out of 3 biological replicates as significantly regulated (Fig. 3A). This resulted in 23 significantly up or down regulated proteins between RTT and iCTR on D3, 111 on D9, 72 on D15 and 243 on D22. We then compared the significantly up or down regulated proteins across different time points in a Venn diagram (**Fig. S3**). We noticed that the majority of the significantly regulated proteins have only 0-2% overlap between the different time points. To better understand the biological processes of the proteins that are differentially expressed at each time point, we performed Gene Ontology (GO) enrichment analysis with respect to biological functions (Fig. 3B). GO analysis on the significant proteins revealed terms related to up regulation in RTT versus iCTR of ‘neuron apoptotic process’ and ‘cellular response to hypoxia’ on D3. Terms related to ‘negative regulation of axon regeneration’, ‘positive regulation of filopodium assembly’ and ‘positive regulation of neuron projection development’ were down regulated in RTT on D3. On D9 of neuronal differentiation, ‘insulin receptor signaling pathway’ was up regulated whereas ‘excitatory postsynaptic potential’ was down regulated in RTT. On D15, ‘cell-cell adhesion’ and ‘acyl-CoA metabolic process’ were up regulated and terms such as ‘axon guidance’, ‘brain development’ and ‘histone acetylation’ were down regulated. Furthermore, on D22, terms related to ‘cell-cell adhesion’ and ‘actin cytoskeleton organization’ were up regulated and ‘nervous system development’ and ‘forebrain development’ were down regulated in RTT. Terms such as brain development were down regulated in D15 as well as D22. To further verify the results of the mass spectrometry, we performed western blot analysis for proteins SOX2 and SOX9, transcription factors with pivotal role in development and differentiation [25, 26], which showed significant differences in expression levels between iCTR and RTT lines at D22 (Fig. 3A, C). In line with mass spectrometry data, western blot analysis showed a significant increase in SOX9 expression levels in RTT lines when compared to iCTR (p=0.0057, unpaired t-test), and a decrease in SOX2 expression in RTT lines at D22, although this did not reach statistical significance (p=0.07, unpaired t-test). Together both approaches demonstrate that SOX2 and SOX9 were differentially expressed in RTT versus iCTR. Overall, we show that proteins associated with neuronal development are differentially expressed in RTT at early stages of neuronal differentiation.

**Figure 3.**
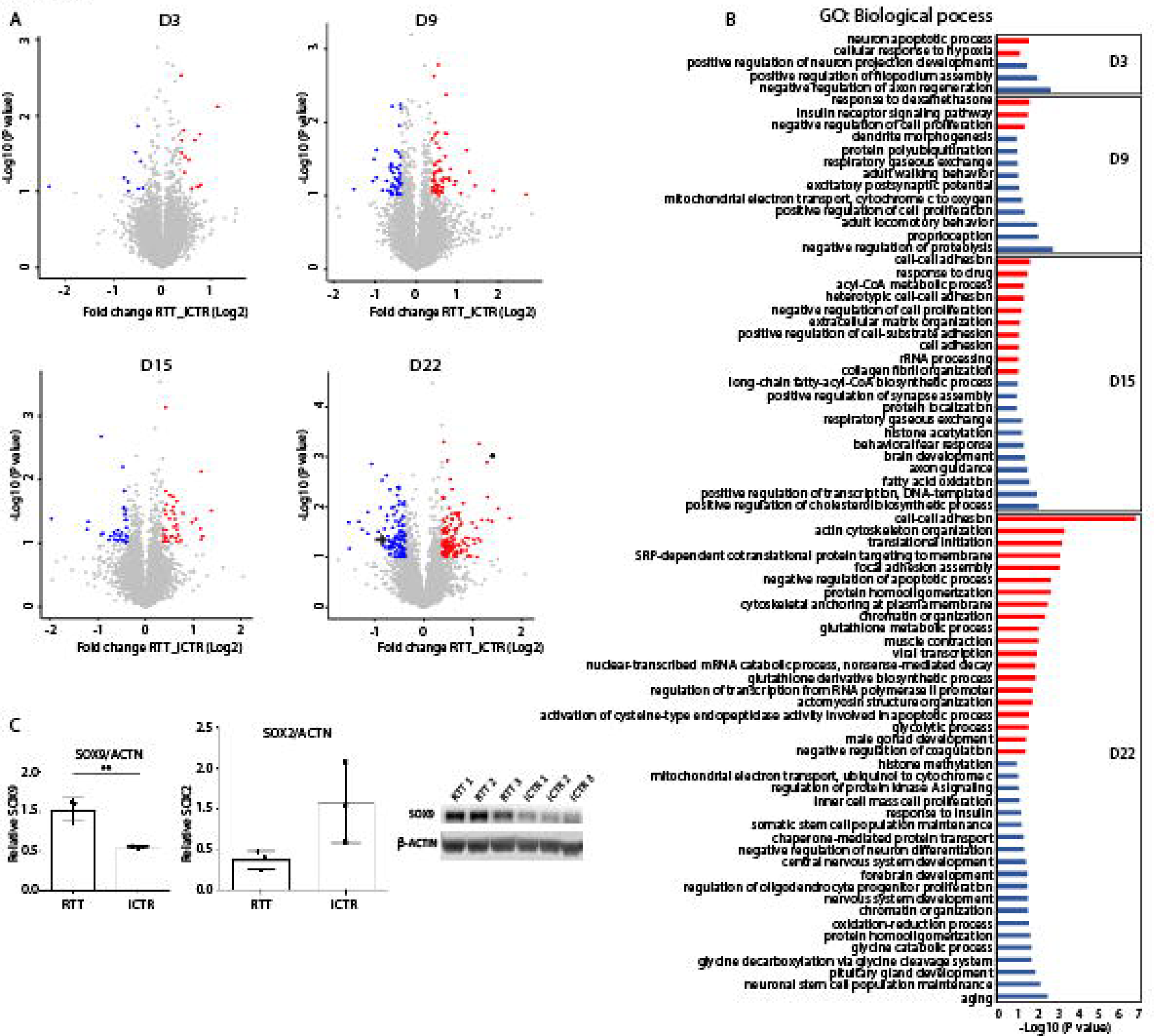
Volcano plots each day and GO analysis. (A) Volcano plot demonstrating proteins differentially regulated in RTT compared to iCTR in each time point of neuronal development. Each data point represents a single quantified protein. The x-axis represents the log2-fold change in abundance (RTT/iCTR) and y-axis the -log10 (p-value). Threshold for significant proteins is chosen for p-value cut-off (0.1) and fold change ≥ 1.3. Proteins in blue indicate for down regulation and in red indicate for up regulation in RTT. Arrow in day 22 points for SOX2 expression and asterisk points for SOX9 expression. (B) Gene Ontology analysis of the significant proteins on their biological function in each time point of neuronal differentiation. Red indicates the up regulated and blue the down regulated biological processes. (C) Western blot analysis showing SOX9 and SOX2 expression in iCTR NES lines at day 22. Significant increase in SOX9 expression in RTT samples and SOX2 shows a trend towards decrease in RTT samples.

### Coordinated proteome alteration during neuronal development in RTT syndrome

To gain insight into how the differentially expressed proteins in RTT behave across time points, we further analysed all the significantly up or down regulated proteins at D3, D9, D15 and D22. This resulted in 234 significantly up and 190 down regulated proteins. The average log2 values of RTT were extracted with iCTR for each time point and the difference between RTT and iCTR is shown in a heat map (Fig 4A). To obtain an unbiased view of the differentially expressed proteins during neuronal differentiation, we performed cluster analysis on the significantly up or down regulated proteins. This resulted in four clusters for both the up or down regulated proteins with distinct expression profiles. Cluster 1 contains proteins strongly up regulated in D3 that are involved in neuron apoptotic processes and cytochrome c release from mitochondria-related GO terms. The majority of the up regulated proteins belong to cluster 2 whose expression levels increase in D22 and are involved in adhesion assembly and glutathione metabolic processes. Cluster 3 contains proteins up regulated in D9 and D15, whereas cluster 4 represents proteins strongly up regulated in D9. In the down regulated proteins, cluster 1 represents proteins that showed a strong down regulation at D15 and that are mainly involved in cholesterol biosynthesis and fatty acid oxidation (Fig. 4B). Cluster 2 represents proteins strongly down regulated in D9, which are associated with regulation of proteolysis and mRNA stability. Cluster 3 of the down regulated proteins in RTT shows a decrease expression profile in D3 which are involved in axon regeneration and neuron projection development and cluster 4 covers many proteins strongly down regulated in D22 involved in processes such as neuronal stem cell maintenance, pituitary gland development and aging. Collectively, our data reveals the expression changes of the differentially expressed proteins during neuronal development in RTT having a stage-specific expression pattern.

**Figure 4.**
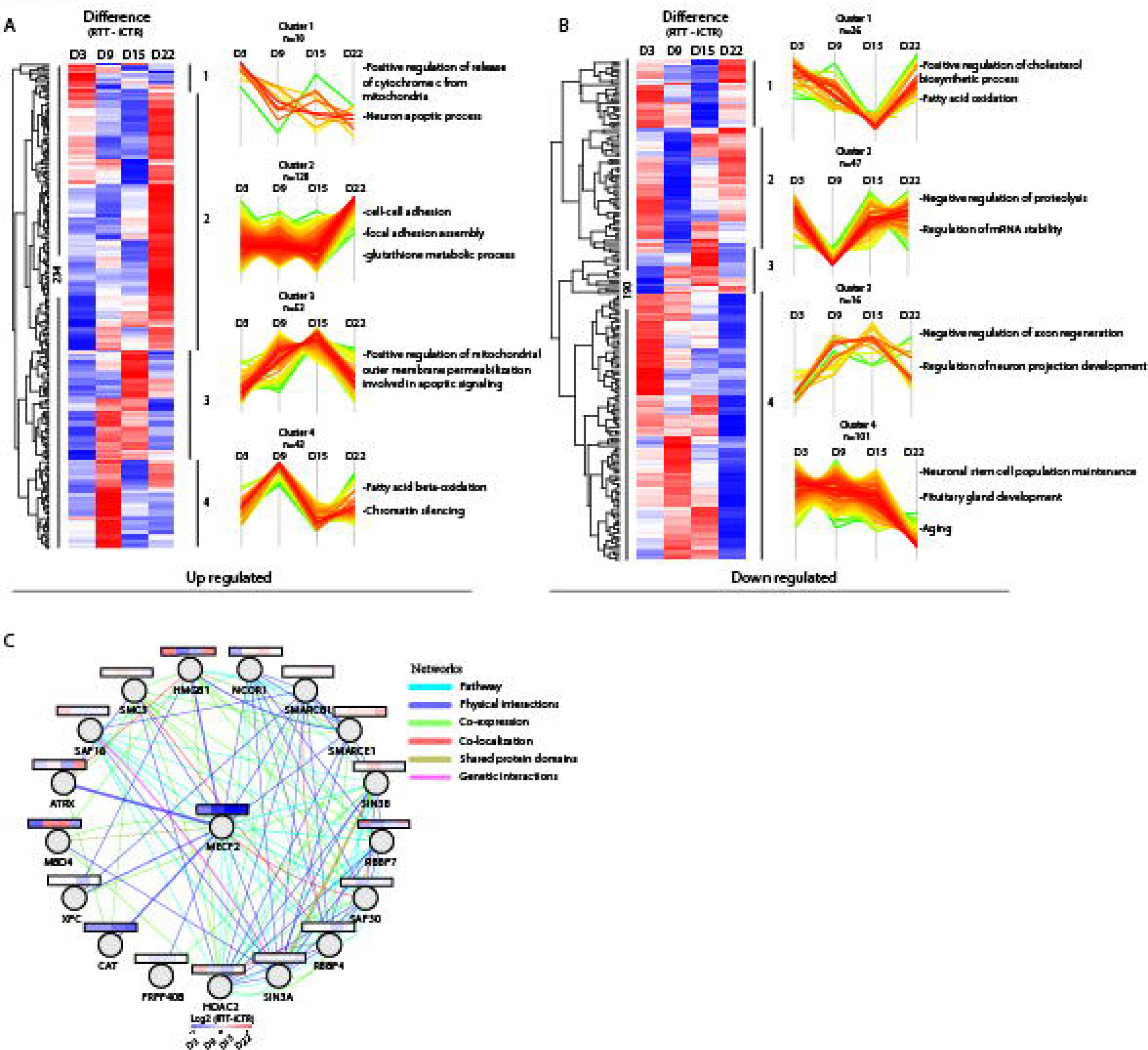
Up and down regulated proteins and MeCP2-binding partners. (A) Heat map of all significant up and (B) down regulated proteins with a p-value ≤ 0.1 and ≥ 1.3 fold change between RTT and iCTR. The Z score of the difference between RTT - iCTR is given for each day with the corresponding cluster analysis and the GO terms for biological processes. (C) Network analysis of MeCP2-binding proteins identified in our data. The average Log2 ratio RTT-iCTR over time is color coded for each protein over the time course of neuronal development. Edges are color coded according to the network as indicated.

### MeCP2 network analysis

To further investigate the proteins that are targets of the MeCP2 protein, we drew a protein interaction network (Cytoscape, Genemania plugin) using MeCP2 protein as input (Fig.4C). The data covered 20 MeCP2-interacting proteins of which 18 proteins were identified in our data. To further investigate how the MeCP2-interacting proteins change over time in RTT, we extracted the average log2 values of RTT from iCTR at each time point. As expected, MeCP2 is down regulated along the course of neuronal differentiation. The proteins are tightly interconnected around HDAC2, SIN3A, RBBP4, SAP30, SMARCE1, SMARCB1, and MECP2, but to a lesser extend around CAT, XPC, PRPF40B, and MBD4. The network revealed several RNA/DNA binding proteins of which MECP2 and CAT are one of the most down regulated proteins in RTT. While some proteins in the network, such as SMARCB1 and SIN3A stayed constant over time, others showed changing levels, such as MBD4 and HMGB1. Interestingly, MBD4 is a member of the methyl-CpG-binding domain (MBD) family of proteins together with MeCP2. Overall, we searched for MeCP2-binding partners and showed how these proteins change along the course of neuronal differentiation in RTT and iCTR.

### Protein subsets differentially expressed at all-time points

To study proteins differentially expressed between RTT and iCTR regardless of the time point of differentiation, we grouped the RTT samples derived from all time points together and iCTR derived from all time points together. Due to the tight ratios typically observed in TMT quantification [27], we selected a cut-off for proteins being up or down regulated with a p-value ≤ 0.1 and ≥ 1.3 fold change difference in RTT compared to iCTR based on the observed distribution in the volcano plot (Fig. 5A). We identified 27 proteins being up and 12 proteins being down regulated in RTT compared to iCTR. As expected, MeCP2 was one of the most strongly down-regulated proteins in RTT. GO analysis revealed biological processes such as ‘cell-cell adhesion’ and ‘acyl-CoA metabolic processes’ to be up regulated, which was up regulated as well in D15 and D22 (Fig. 4A, 5B). In contrast, several processes such as ‘response to cadmium ion’, ‘response to drug’ and ‘behavioral fear response’ were down regulated in RTT (Fig. 5B). Analysis of the differentially regulated proteins using Reactome pathway analysis revealed among others, ‘JAK/STAT signaling after Interleukin-12 stimulation’ and ‘regulation of MeCP2 expression and activity’ to be differentially expressed in RTT versus iCTR (Fig. 5C). To further visualize the connectivity among these significant proteins, we analyzed their protein networks in the Cytoscape tool (Genemania plugin). A high degree of connectivity, such as being co-expressed and having shared genetic interactions, around these proteins was identified. Interestingly, the majority of the proteins are involved in immunity, actin cytoskeleton organization and calcium binding (Fig. 5D). Together, we show that proteins associated with immunity and metabolic processes are differentially expressed in RTT in a time-point independent manner during differentiation towards neurons.

**Figure 5.**
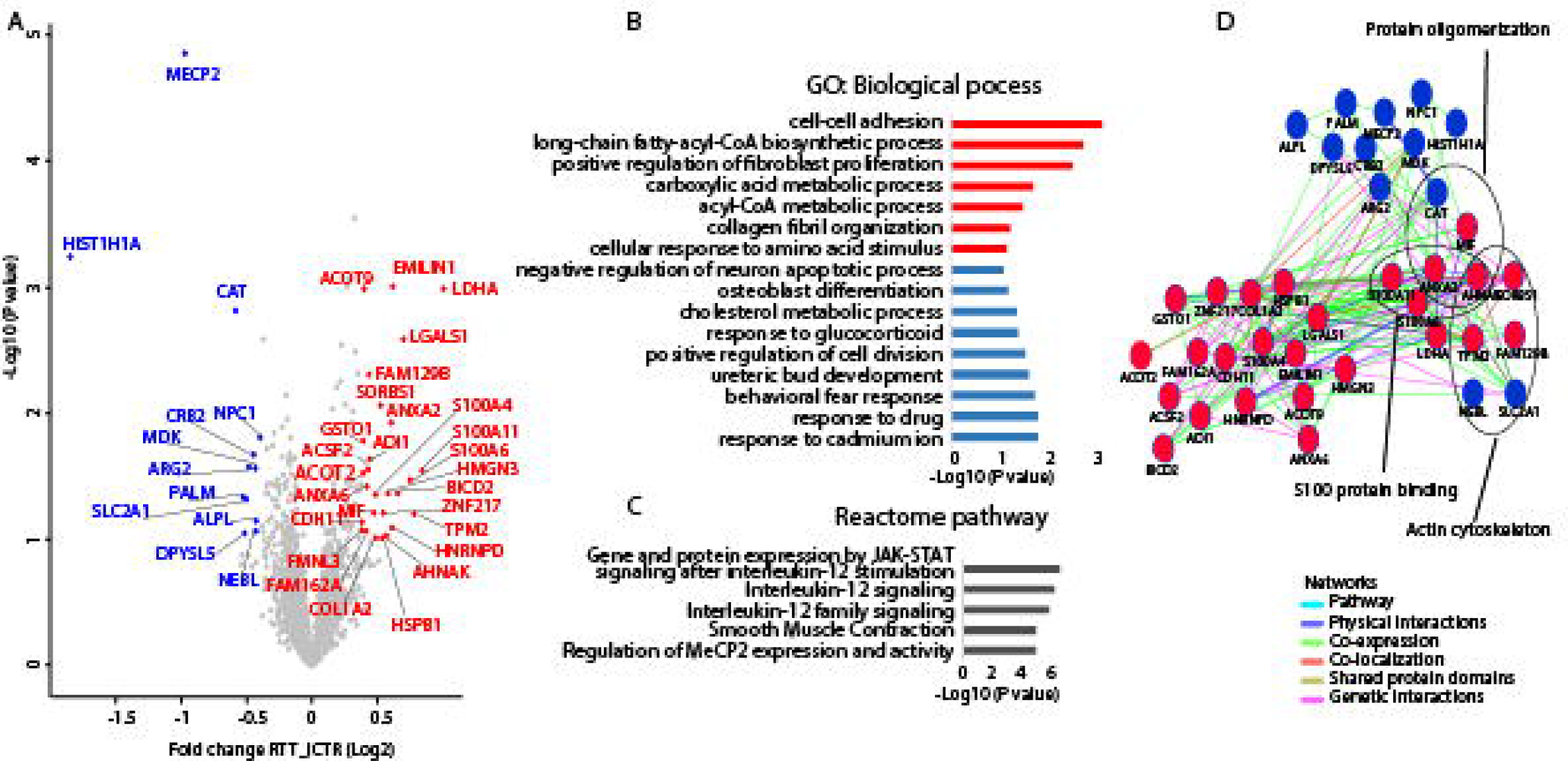
Volcano plot all days and network analysis. (A) Volcano plot demonstrating proteins differentially expressed in RTT versus iCTR after pooling all time points of neuronal development. The x-axis represents the log2 fold change in abundance (RTT/iCTR) and y-axis the -log10 (p-value). Threshold for significant proteins is chosen for p-value cutt-off (0.1) and fold change ≤ 1.3. Up regulated proteins in RTT are shown in red and down regulated are shown in blue. (B) GO analysis on the Biological Process of the significant proteins. X-axis represents the -Log10 (p-value) and red and blue colors indicate for up and down regulated proteins in RTT respectively (C) Reactome pathway analysis of the significant proteins (D) Network analysis of the significant proteins by Cytoscape plugin GeneMania. Red indicates for up regulated and blue indicates for down regulated proteins in RTT.

## Discussion

Before typical RTT-associated symptoms appear, both RTT patients and mouse models of RTT already show abnormalities [28, 29]. This implies that underlying mechanisms are already affected during early neurodevelopmental stages. Much of our understanding of how MeCP2 deficiency contributes to RTT disease is derived from genomic and transcriptomic studies. So far, only a few proteomic studies have been performed on RTT human derived tissue [19, 30, 31]. The current study provides mass spectrometry-based quantitative proteomic data, depth of about 7000 proteins, using an earlier developed iPSC-based models involving RTT patient cells and isogenic controls [24]. We showed that changes in dendrite morphology or synaptic defects, previously associated with RTT [22, 32], already become apparent at early developmental stages. Proteins involved in immunity and metabolism, also in line with previous studies on RTT pathology [33, 34], are differentially expressed at all time points. This indicates we found time-point specific alterations as well as differentially expressed proteins at all time points during early neuronal differentiation in RTT cultures. Insight into differentially expressed protein levels could support identification of novel biomarkers as well as therapeutic strategies.

### RTT samples present protein expression changes that became more apparent from early to late neuronal stem cell phases

Already at D3 of neural induction, we found decreased protein levels associated with axon regeneration and filopodium assembly, and increased protein levels in neuronal apoptotic processes. Furthermore we show a down regulation of proteins associated with dendrite morphogenesis and excitatory postsynaptic potential at D9, and down regulation of axon guidance and brain development at D15 of neural induction. By D22 a clear set of proteins was altered in RTT with up regulated proteins associated with ‘cell adhesion’, ‘cytoskeleton organization’ and ‘translation initiation’. Proteins that are down regulated in RTT are involved in ‘nervous system development’, ‘forebrain development’ as well as ‘histone methylation’, which has classical roles with MeCP2 in its organization [35]. While the proteomic alterations found are relatively small (1.3 fold-change), they are in line with previous studies showing dysregulation of dendrites and axons in RTT patients/mouse models [33, 36-38] as well as in the juvenile RTT brain [39-41]. Although at these time points of neuronal stem cells did not develop dendrites or axons yet, our findings indicate that proteins involved in these processes are already expressed and altered in RTT at early developmental stages. During day 3-22, the number of proteins that are differentially expressed increases, although this set of protein changes over the time points. The increase of the fold change over time is an indicator of the manifestation of RTT, starting from early brain development towards the mature central nerve system. Furthermore, we noticed a small overlap of the altered proteins across time points which might indicate that proteins associated with RTT are specifically expressed at distinct time-points during the course of neural differentiation and reflect the robust cellular identity changes during early developmental stages. Overall, we show that dysregulation of MeCP2 affects protein expression changes associated with neurodevelopmental functions at early stages of neuronal differentiation.

### Expression changes of MeCP2-interacting proteins in RTT and iCTR

Network analysis using GeneMANIA [42] revealed part of these proteins to be interacting with MeCP2. Interestingly, of these interacting proteins, MBD4 is a member of the methyl-CpG-binding domain (MBD) family of proteins together with MeCP2. Mutations in these functional important domains tend to cause RTT associated phenotypes [43-45]. Furthermore, HMGB1, which was down regulated in D9 and D15 in our data, was previously shown to be lower expressed in hippocampal granule neurons of MECP2 KO mice [46].

### Robust protein profile changes along the course of neuronal differentiation

When all RTT versus iCTR samples from different time points were pooled, data revealed differentially expressed proteins in RTT involved in immunity, calcium binding and metabolism. This analysis allowed us to compensate for limited amount of biological replicates in such high-throughput technology. Several proteins associated in metabolic processes were differentially expressed in RTT, including ACSF2, ACOT2, ACOT9, LDHA, MIF, NPC1, CAT and GSTO1. Current evidence in perturbed lipid metabolism in the brain and the peripheral tissue of RTT patients and mouse models now also supports the metabolic dysfunctions as a component of RTT [34]. Also mRNA GSTO1 was up regulated in RTT patients’ lymphocytes together with several other mitochondrial related genes [47]. Furthermore, our results indicate dysregulated proteins associated with Interleuking-12 signalling. Interestingly, children with MeCP2 duplication show immunological abnormalities and suppressed IFN-□ [48]. Although RTT is more often classified as a neurodevelopmental disorder, recent studies in cytokine release also suggest involvement of the immune system [49]. We further identified several proteins associated with calcium signalling to be altered. A disturbance in calcium homeostasis during early postnatal development was reported in MeCP2 knockout model and altered calcium signalling RTT-iPSC-derived cells [50, 51]. Altogether, this indicates that next to dysregulation in neurodevelopmental processes, disease mechanisms underlying RTT phenotypes could also involve immunity, calcium signaling and metabolism.

### Early treatment

The finding, that neuronal stem cells of RTT patients do show altered protein expressions responsible for neuronal development and maturation, indicates that RTT influences the patients much earlier than first symptoms actually appear. Therefore, possibilities for early treatment need to be discussed. As there is no cure for RTT yet, mutations on MeCP2 are not screened for in pre-natal diagnostics. However, an early testing post-natal could make sense, as we did show that the disease has its onset already before symptoms appear. The time window after birth until approximately 18 month could therefore be used to support neuronal stimulation and maturation with treatments such as IGF-1 or Bumetanide [52-54]. This could support or extend the compensatory process patients’ show before the start of symptoms and maybe extenuate the disease onset. The progress of RTT is based on different aspects. There is evidence that mosaicism and therefore the ratio between cells expressing healthy compared to mutated MeCP2 plays a major role [55]. Furthermore, the mutation itself has a huge impact on severity of the disease [56]. However, in almost all cases of patients with MeCP2 mutation a post-natal phase of compensation can be observed. We are convinced that this critical period can be used to start an efficient treatment and prevent disease progression. Therefore, we hope that our results give awareness of the early pre-natal onset of RTT and emphasize the need of early testing followed by an effective treatment.

## Experimental procedures

### Cell culture and isogenic controls

RTT patient fibroblasts were derived from the *Cell lines and DNA bank of Rett syndrome, X-linked mental retardation and other genetic diseases* at the University Siena in Italy via the Network of Genetic Biobanks Telethon. Three fibroblast lines carrying different MECP2 mutations were selected. RTT#2282C2 showing a deletion in Exon 3 and 4 of the MeCP2 gene (RTT Ex3-4), RTT#2204 carrying a missense mutation within the methyl binding domain (RTT T158M) and RTT#2238 having a nonsense mutation in the transcription repression domain (RTT R255X). Fibroblasts were derived frozen and thawed and expanded in in fibroblast medium (DMEM-F12, 20% FBS, 1%NEAA, 1%Pen/Strep, 50 μM β-Mercaptoethanol). To generate pure RTT fibroblast lines and isogenic controls cells were detached from cell culture plate and single fibroblasts were seeded in wells of a 96-well plate. Cells were further expanded and characterized on their MeCP2 state by immunocytochemistry and PCR [24]. All of our experiments were exempt from the approval of the institutional review board.

### Reprogramming

Reprogramming of fibroblasts was performed as described before [24]. In brief, fibroblasts were detached from cell culture plate and washed with PBS. 4×10^5^ cells were resuspended in 400 µl Gene Pulser^®^ Electroporation Buffer Reagent (BioRad) with 23,4 µg of each episomal plasmid (Addgene, Plasdmid #27078, #27080, #27076) containing the reprogramming factors OCT4, SOX2, KLF4 and C-MYC. Cell solution was carefully mixed and electroporated with three pulses of 1.6 kV, capacitance of 3 μF and a resistance of 400 Ω (Gene Pulser II (BioRad)). Fibroblasts were left for recovery in Fibroblast medium without antibiotics containing 10 µM Rock inhibitor (Y-27632). After cells reached a confluence of 60-70%, medium was changed to TeSR™-E7™ (STEMCELL). Colonies appeared after 21-28 days. They were picked manually and maintained in TeSR™-E8™ (STEMCELL).

### Differentiation of neuronal stem cells

In total, 8 iPSC lines were differentiated towards neuronal stem cells as described before [21, 57]. Four RTT lines and four isogenic controls from the same individual therefore, were plated in high-density on Geltrex^®^-coated wells of a 12-well plate in TeSR™-E8™ with 10 µM Rock inhibitor. Medium was changed daily for 2 days. Afterwards half of the medium was changed daily to Neuro-Maintenance-Medium (NMM) (1:1 DMEM/F12+GlutaMAX:Neurobasal Medium, 1x B27, 1xN2, 2,5 µg/ml Insulin, 1,5 mM L-Glutamin, 100 µM NEAA, 50 µM 2-Mercaptoethanol, 1% penicillin/streptomycin) containing 1 µM Dorsomorphin and 10 µM SB431542 up to day 12. At day 10-12 rosette structures appeared, which were manually picked and further cultured on Poly-L-Ornithin (0,01%)/Laminin (20 µg/ml) coated cell culture plates in NMM medium containing EGF (20 ng/ml) and FGF-2 (20 ng/ml). Half of medium was changed daily and cells were cultured up to day 25.

### Immunocytochemistry

To perform immunocytochemistry cells were fixated with 4% Paraformaldehyde and blocked with blocking buffer containing 5% Normal Goat Serum (Gibco^®^), 0,1% bovine serum albumin (SigmaAldrich) and 0,3% Triton X-100 (SigmaAldrich). Primary antibody incubation for MeCP2 (D4F3, CellSignaling, 1:200, rabbit), OCT3/4 (C-10, Santa Cruz, 1:1000, mouse), SSEA4 (Developmental Studies Hybridoma Bank, 1:50, mouse), TRA1-60 (Santa Cruz, 1:200, mouse), TRA1-81 (Millipore, 1:250, mouse), SOX2 (Millipore, 1:1000, rabbit) was performed in blocking buffer over night at 4°C. Next day cells were washed and secondary antibody Alexa Fluor^®^ 488 (ThermoFisher, 1:1000, mouse or rabbit) and Alexa Fluor^®^ 594 (ThermoFisher, 1:1000, mouse or rabbit) were applied in blocking buffer for 1 h at room temperature. To identify cell nuclei DAPI was used for 5 min before cells were mounted with Fluoromount™ (Sigma-Aldrich).

### RNA collection, Sequencing and PCR analysis

To isolate RNA samples, standard TRIzol^®^-Chloroform isolation was done. RNA was stored at −80°C until further processed. For RNA-Seq, RNA-quality was validated, using Agilent 2200 TapeStation system. When samples showed RNA Integrity Number (RIN) > 8 they were collected and RNA was further processed according to manufacturer’s protocol (Illumina, Catalog # RS-122-9004DOC) followed by sequencing by Illumina HiSeq 4000 (50 base pair single read).

For PCR analysis RT-PCR was performed. cDNA was synthesized by using SuperScriptIV-Kit (ThermoFisher) following manufacturer’s recommendations and could be stored until further processing at −20°C. To perform PCR different primer sets were used (Tab.1) and PCR was executed with Phire Hot Start II DNA Polymerase (ThermoFisher).

### Western blotting

Frozen cell pellets were lysed by adding WB-Lysate buffer (50mM Hepes ph 7,5, 150 mM NaCl, 1 mM EDTA, 2,5 mM EGTA, 0,1% TritonX-100, 10% Glycerol, 1 mM DTT). To determine protein concentration Bradford-Test was performed and 30 ug of sample were used. For SDS-PAGE pre-casted Gels were used (Biorad) and ran in 10× Tris/Glycine Buffer for Western Blots and Native Gels (Biorad #1610734). Gels were blotted in tank-blotter (Biorad) on PVDF membranes (Biorad) according to manufactures protocol. After protein transfer blots were blocked in 5% BSA/TBS for 1 h and stained for Sox2 (1:100, Millipore AB5603), Sox9 (1:250; CellSignalling 82630) and ß-actin (1:1000; Chemicon, C4 MAB 1501) in 5% BSA/TBS over night at 4°C. Next day blots were washed and stained with secondary antibodies in 5% BSA/TBS for 1 h at RT. After another 3 TBS washes blots were stained with SuperSignal™ West Femto Maximum Sensitivity Substrate (ThermoFisher) and analysed with LiCor analyser.

### Sample collection

Samples were collected at different days throughout the differentiation. First samples were taken at day 3 of protocol, one day after medium change towards NMM with Dorsomorphin and SB431542. Second samples were taken at day 9, before rosette structures were cut, followed by third sample collection at day 15, after rosettes were manually picked. Finally, fourth samples were taken at day 22, after first passage was performed and cells were recovered. To collect all, cells were washed once with PBS and then scraped off the cell culture plate. Solution was collected in an Eppendorf Microtube and centrifuged at maximum speed for 5 min. Supernatant was discarded and pellet was frozen at −80°C until further processed for mass spectrometry.

### Cell lysis and protein digestion

Samples were lysed, reduced and alkylated in lysis buffer (1 % sodiumdeoxycholate (SDC), 10 mM tris(2-carboxyethyl)phosphine hydrochloride (TCEP), 40 mM chloroacetamide (CAA) and 100 mM TRIS, pH 8.0 supplemented with phosphatase inhibitor (PhosSTOP, Roche) and protease inhibitor (Complete mini EDTA-free, Roche). After sonication, samples were centifugated at 20,000 × g for 20 min. Protein concentration was estimated by a BCA protein assay. Reduction was done with 5 mM Ammonium bicarbonate and dithiothreitol (DTT) at 55°C for 30 min followed by alkylation with 10 mM Iodoacetamide for 30 min in dark. Proteins were then digested into peptides by LysC (Protein-enzyme ratio 1:50) at 37°C for 4 h and trypsin (Protein-enzyme ratio 1:50) at 37°C for 16 h. Peptides were then desalted using C18 solid phase extraction cartridges (Waters).

### Tandem Mass Tag (TMT) 10 plex labelling

Aliquots of ~ 100 µg of each sample were chemically labeled with TMT reagents (Thermo Fisher) according to Figure 1. In total three TMT mixtures were created for each biological replicate. Peptides were resuspended in 80 µl resuspension buffer containing 50 mM HEPES buffer and 12.5 % acetonitrile (ACN, pH 8.5). TMT reagents (0.8 mg) were dissolved in 80 µl anhydrous ACN of which 20 µl was added to the peptides. Following incubation at room temperature for 1 hour, the reaction was then quenched using 5% hydroxylamine in HEPES buffer for 15 min at room temperature. The TMT-labeled samples were pooled at 1:1 ratios followed by vacuum centrifuge to near dryness and desalting using Sep-Pak C18 cartridges.

### Off-line basic pH fractionation

Before the mass spectrometry analysis, the TMT mixture was fractionated and pooled using basic pH Reverse Phase HPLC. Samples were solubilized in buffer A (5% ACN, 10 mM ammonium bicarbonate, pH 8.0) and subjected to a 50 min linear gradient from 18 % to 45 % ACN in 10 mM ammonium bicarbonate pH 8 at flow rate of 0.8 ml/min. We used an Agilent 1100 pump equipped with a degasser and a photodiode array (PDA) detector and Agilent 300 Extend C18 column (5 μm particles, 4.6 mm i.d., and 20 cm in length). The peptide mixture was fractionated into 96 fractions and consolidated into 24. Samples were acidified with 10% formic acid and vacuum-dried followed by re-dissolving with 5% formic acid/5% ACN for LC-MS/MS processing.

### Mass spectrometry analysis

We used nanoflow LC-MS/MS using Orbitrap Lumos (Thermo Fisher Scientific) coupled to an Agilent 1290 HPLC system (Agilent Technologies). Trap column of 20 mm × 100 µm inner diameter (ReproSil C18, Dr Maisch GmbH, Ammerbuch, Germany) was used followed by a 40 cm × 50 µm inner diameter analytical column (ReproSil Pur C18-AQ (Dr Maisch GmbH, Ammerbuch, Germany). Both columns were packed in-house. Trapping was done at 5 µl/min in 0.1 M acetic acid in H2O for 10 min and the analytical separation was done at 100 nl/min for 2 h by increasing the concentration of 0.1 M acetic acid in 80% acetonitrile (*v/v*). The mass spectrometer was operated in a data-dependent mode, automatically switching between MS and MS/MS. Full-scan MS spectra were acquired in the Orbitrap from m/z 350-1500 with a resolution of 60,000 FHMW, automatic gain control (AGC) target of 200,000 and maximum injection time of 50 ms. Ten most intense precursors at a threshold above 5,000 were selected with an isolation window of 1.2 Da after accumulation to a target value of 30,000 (maximum injection time was 115 ms). Fragmentation was carried out using higher-energy collisional dissociation (HCD) with collision energy of 38% and activation time of 0.1 ms. Fragment ion analysis was performed on Orbitrap with resolution of 60,000 FHMW and a low mass cut-off setting of 120 m/z.

### Data processing

Mass spectra were processed using Proteome Discover (version 2.1, Thermo Scientific). Peak list was searched using Swissprot database (version 2014_08) with the search engine Sequest HT. The following parameters were used. Trypsin was specified as enzyme and up to two missed cleavages were allowed. Taxonomy was set for Homo sapiens and precursor mass tolerance was set to 50 p.p.m. with 0.05 Da fragment ion tolerance. TMT tags on lysine residues and peptide N termini and oxidation of methionine residues were set as dynamic modifications, and carbamidomethylation on cysteine residues was set as static modification. For the reporter ion quantification, integration tolerance was set to 20 ppm with the most confident centroid method. Results were filtered to a false discovery rate (FDR) below 1%. Finally, peptides lower than 6 amino-acid residues were discarded. Within each TMT experiment, reporter intensity values were normalized by summing the values across all peptides in each channel and then corrected for each channel by having the same summed value. After that the normalized S/N values were summed for all peptides. Finally proteins were Log_2_ transformed and normalized by median subtraction.

### Data visualization

The software Perseus was used for data analysis and to generate the plots. Volcano plots for each time point was generated and up- or down-regulated proteins were considered significant with a fold change cut-off = 1.3. Functional analysis to enrich to GO terms were done on David Database and pathway enrichment analysis was done on Reactome Functional Interaction (http://www.reactome.org/). Furthermore, protein interaction network was performed using Cytoscape, Gnenmania plugin.

### Data availability

All mass spectrometry proteomics data have been deposited to the ProteomeXchange Consortium via the PRIDE partner repository with the dataset identifier PXD013327

## Acknowledgements

Specimens were provided by the *Cell lines and DNA bank of Rett Syndrome, X-linked mental retardation and other genetic disease*, member of the Telethon Network of Genetic Biobanks (project no. GTB12001), funded by Telethon Italy, and of the EuroBioBank network.

---

**Figure S1. iPSC characterisation.** (A) Exemplary characterisation of iPSC lines. Immunocytochemistry for pluripotency marker (OCT3/4, SOX2, SSEA4, TRA1-60, TRA1-81). (B) PCR-analysis for pluripotency marker. (C) PluriTest plot for RTT and iCTR iPSC lines (yellow).

**Figure S2. Number of proteins identified.** A bar chart showing the number of proteins identified in each biological replicate and time point when analyzed as 20× high-pH fractions.

**Figure S3. Venn diagram.** (A) Number of proteins decreased in expression in RTT at different time points. (B) Number of proteins increased in expression in RTT at different time points. (C) Overview of the number of proteins altered in RTT.

