## Supplementary figures and images for "Quantitative proteomic alterations of human iPSC-based neuronal development indicate early onset of Rett syndrome"

### figure S1

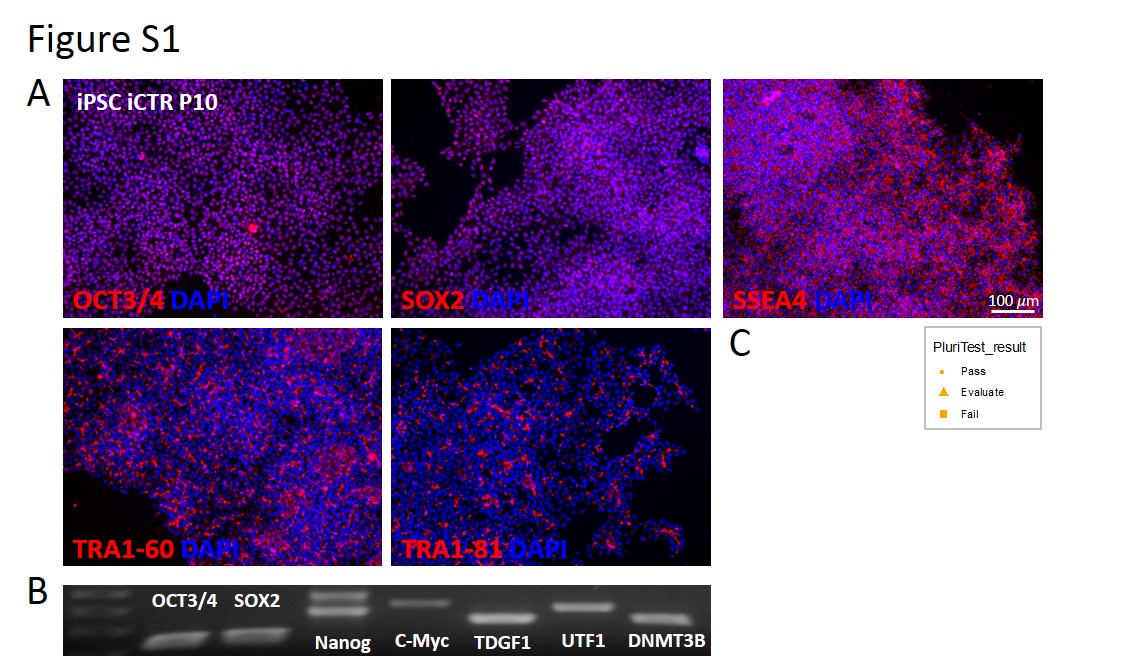

### figure s2

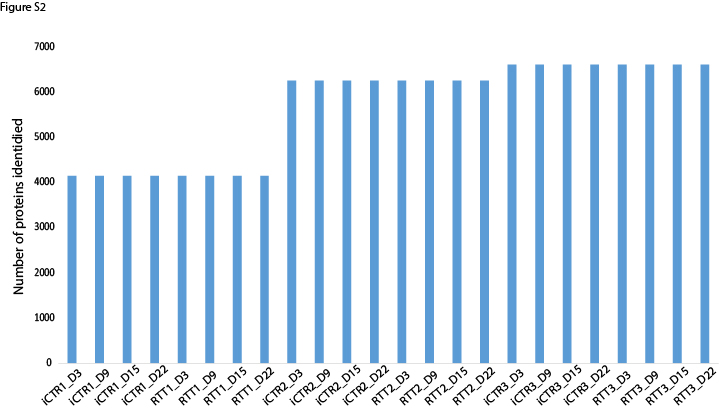

### figure s3

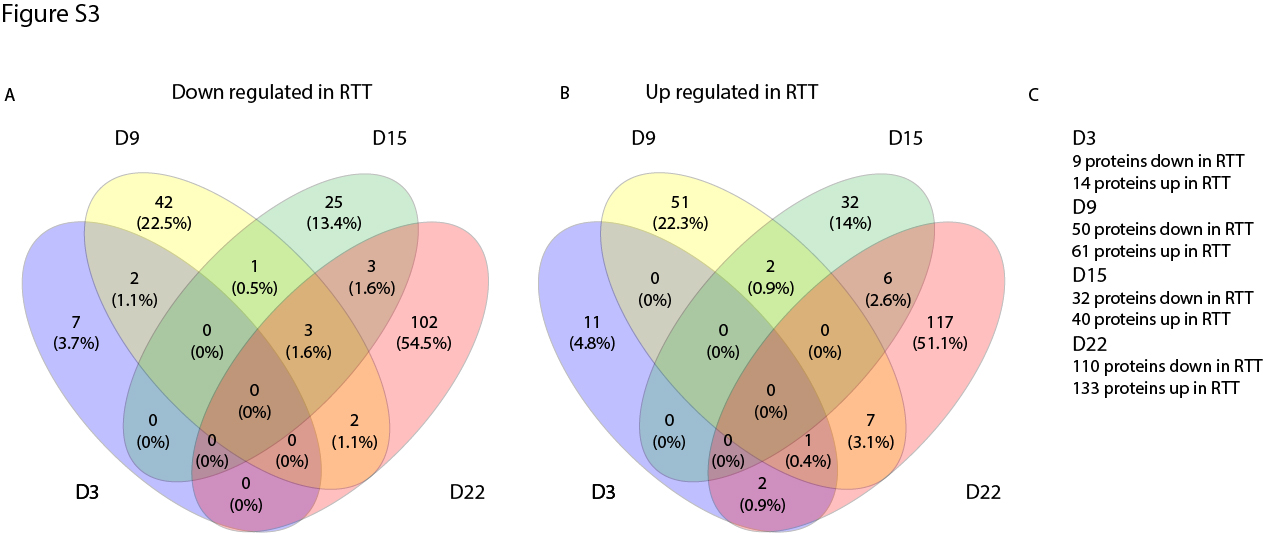
